# Functional gene analysis and cultivation experiments predict the degradation of diverse extracellular polysaccharides by ubiquitous taxa in pustular microbial mats from Shark Bay, Western Australia

**DOI:** 10.1101/2022.05.18.492586

**Authors:** Elise Cutts, Clemens Schauberger, Emilie Skoog, Tanja Bosak

## Abstract

Microbial exopolymeric substances (EPSs) form the organic, polysaccharide-rich matrix of marine microbial mats and can mediate the binding and precipitation of carbonate minerals therein. Here, we investigate the molecular ecology of carbohydrate degradation and production in pustular mats from Shark Bay, Western Australia, by analyzing 84 metagenome-assembled genomes (MAGs) and the composition of microbial communities enriched from a pustular mat on various polysaccharide substrates. The annotation of 4000 genes from hundreds of carbohydrate-active enzyme (CAZyme) families in the MAGs and mapping of polysaccharide-degrading CAZymes to their predicted substrates identify trends in the distribution and localization of degradation-associated CAZymes across different bacterial phyla. The compositions of microbial communities enriched on a range of polysaccharides inoculated with pustular mat material support the predicted trends. The combined metagenomic and experimental analyses reveal a widespread potential for EPS degradation among MAGs from Shark Bay pustular mats and suggest distinct roles for some phyla that are reported at high abundances in mats. Specifically, Bacteroidetes are likely to be primary degraders of polysaccharide EPSs, alongside Planctomycetes and a small subset of Alphaproteobacteria and Gammaproteobacteria. Planctomycetes, some Bacteroidetes, Verrucomicrobia, Myxococcota and Anaerolineae are also predicted to favor degradation of sulfated substrates, which are present in the EPS matrix of pustular mats. Large sets of functionally varied CAZymes without signal peptides tagging them for export implicate Anaerolineae and Verrucomicrobia in degrading the downstream products of primary EPS degradation.

**Importance:** Modern marine microbial mats are rich in exopolymeric substances (EPSs) — complex, high molecular weight polymers secreted by bacteria — that mediate the formation of carbonate minerals and the preservation of microbial textures in mats. However, the organisms involved in EPS cycling in these mats have not been identified and the links between EPS degradation, carbonate precipitation, and microbial ecology in mats remain poorly understood. We define distinct roles in EPS cycling for many major microbial taxa that are both ubiquitous and abundant in pustular microbial mats from Shark Bay, Australia. The large genomic potential of these microbes for the modification and degradation of diverse extracellular organic polymers provides a blueprint for future studies aimed at quantifying and verifying the specific contributions of these microbes to EPS degradation, carbon cycling and carbonate precipitation.

## Introduction

Microbial mats are benthic communities of microorganisms embedded within a sticky, thick matrix of biopolymer secretions. These secretions, or exopolymeric substances (EPSs), typically account for ∼90% of biofilm dry mass and, in cyanobacterial mats, are mostly polysaccharides (1–4). The EPS matrix is simultaneously the physical environment for and a biochemical and metabolic extension of the mat community. Anionic EPSs bind cells, certain nutrients (e.g., cations), and excreted digestive enzymes, giving mats a “three-dimensional architecture” that allows organisms to interact and carry out extracellular activities with spatial precision unachievable in a free-living state (5). In marine microbial mats, the matrix also interacts with minerals that are either captured from the environment (i.e., “trapped and bound”) or precipitated in-place. Some mats lithify extensively, forming the morphologically and texturally diverse structures collectively termed microbialites (6). EPSs can influence mineralization in microbialites by providing mineral nucleation sites, inhibiting nucleation (7, 8) and binding or releasing cations such as calcium or magnesium (1, 9–11). The EPS matrix may also indirectly influence the alkalinity engine by generating organic substrates for microbial metabolisms that increase or decrease carbonate alkalinity (6, 10). EPSs thus act as an interface between geochemical processes in mats and their microbial ecology.

The EPS-rich hypersaline pustular mats of Hamelin Pool Shark Bay, Australia (Fig. 1) are the contemporary analogues to 2-billion-year-old peritidal microbial mats that preserve the oldest diagnostic cyanobacterial fossils (12). A relatively consistent picture of the microbial communities in these mats has emerged from 16S rRNA amplicon, metagenomic and metatranscriptomic sequencing, with Proteobacteria (Alpha, Gamma, and Delta), Bacteroidetes, Planctomycetes, Cyanobacteria, Actinobacteria, and Chloroflexi (mainly Anaerolineae) as the most abundant taxa. Cyanobacteria, the most important primary producers in pustular mats, have a relative abundance of ∼5-10% and are mostly active in the upper layers of mats (13–16). Bacteroidetes, Alphaproteobacteria and Cyanobacteria group together in the upper layers, where they are hypothesized to form “phototrophic consortia” (14). Verrucomicrobia are typically observed at ≤3% relative abundance (13, 14, 16). Spirochaetes, Caldithrix, Acidobacteria, Firmicutes, Gemmatimonadetes, and a number of uncultured organisms are typically present in lower abundances of 1–2% (13, 14, 16).

**Figure 1.**
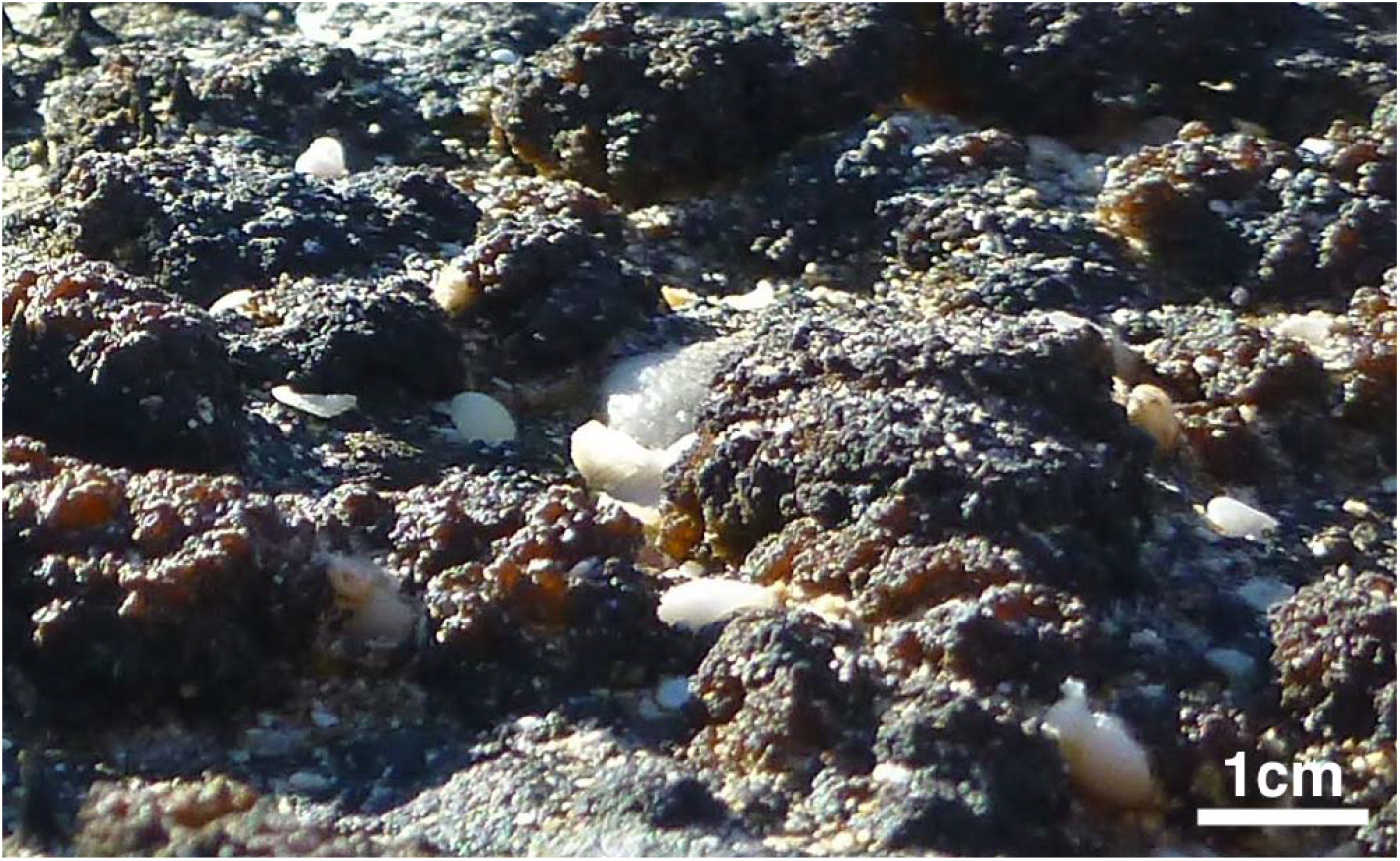
Pustular mats of Hamelin Pool, Shark Bay, Australia. Peritidal pustular mats of Hamelin Pool are centimeter-scale benthic biofilms rich in exopolymeric substances. The photograph shows exposed brown and black mats with millimeter-scale pustules. White grains around some mats are shell fragments.

EPSs are rapidly produced and degraded in mats. These processes change the acidity of EPSs (9, 10) and leave recalcitrant, high molecular weight (HMW) remnants that resist further degradation (10, 17). The degradation of components of the EPS matrix is thought to provide organic carbon to the microbes and control the cycling of oxygen, carbon, and calcium in the mats. These processes require carbohydrate-active enzymes (CAZymes) such as glucosidases and galactosidases, which are active in the slurries of biomineralizing mats from Eleuthera Island, The Bahamas (10). Yet, the links between EPS degradation, carbonate precipitation, and microbial ecology in microbial mats remain poorly understood. Pustule-forming cyanobacteria, embedded in thick, multilayered envelopes, are the primary sources of sulfated and other EPS in Shark Bay pustular mats (18, 19), but most organisms capable of degrading complex polysaccharides in EPS in these and other marine mats have not been identified. Bacteroidetes have been speculated to degrade HMW organic carbon in Shark Bay based on their role in other systems (14, 15), but to our knowledge, their potential and actual contributions to polysaccharide degradation in hypersaline mats are yet to be demonstrated. CAZymes are enriched in the metagenomes from the surficial, EPS-rich layers of pustular mats, where glycoside hydrolase family GH13, glycosyltransferase family GT4, and glycosyltransferase family GT2 were identified as the most abundant CAZyme families (20). However, specific microbial groups that contain these genes, the potential substrates of enzymes encoded by these genes, and potential implications for mat ecology and EPS cycling remain unknown.

We explored the polysaccharide degradation potential in pustular microbial mats from Shark Bay, Australia, by analyzing 84 medium-to-high quality metagenome-assembled genomes (MAGs) from the rehydrated mats and enriching microbes on specific predicted substrates. High potential for polysaccharide synthesis in the Cyanobacteria MAGs is consistent with previous observations identifying Cyanobacteria as the major source of EPS in mats. Our analyses of over 4000 predicted CAZyme genes in this metagenome reveal a taxonomically widespread carbohydrate utilization potential, suggest that Bacteroidetes and Planctomycetes together with a small subset of Alphaproteobacteria and Gammaproteobacteria are primary degraders of extracellular polysaccharides. The same groups and Anaerolineae, Myxococcota, Verrucomicrobia, Hydrogenedentes are also predicted to specifically degrade sulfated extracellular polysaccharides. Large sets of functionally varied CAZymes without signal peptides tagging them for export implicate Anaerolineae and Verrucomicrobia in “downstream” degradation of carbohydrates. 16S rRNA amplicons of enrichment cultures established on a set of polysaccharides support the predicted roles of Bacteroidetes as putative polysaccharide degraders adept at processing complex cyanobacterial EPS, including sulfated EPS, Planctomycetes as specialist degraders of sulfated polysaccharides and Alphaproteobacteria and Gammaproteobacteria as organisms that grow well on specific non-sulfated polysaccharides.

## Results

### CAZymes in the metagenome of pustular mats from Shark Bay

Table 1 summarizes the distributions of the major CAZyme categories — glycoside hydrolase (GH), polysaccharide lyase (PL), carbohydrate esterase (CE), glycosyl transferase (GT), carbohydrate-binding module (CBM), and accessory activity (AA) — and the abundances of genes and gene families within those groups. Analyses of these genes consider both the counts of *unique* CAZymes genes contained within each MAG and the numbers of different CAZyme families represented by these genes. The advantages of and caveats to this approach are addressed in the discussion. Enzymes with predicted activities on carbohydrate substrates were numerous and widely distributed across the 84 MAGs of Shark Bay pustular mats. Broadly speaking, GH, PL, and CE are associated with degradation of carbohydrates, although some GH act as branching enzymes in the synthesis of branched polysaccharides (21), whereas GT, which catalyze the formation of glycosidic bonds, are associated with carbohydrate synthesis.

**Table 1.** Summary of major CAZyme classes in a Shark Bay pustular mat metagenome. Basic summary of the distribution of the major categories of CAZymes — GH, PL, CE, GT, CBM, and AA — in the Shark Bay pustular mat metagenome. Counts of unique CAZymes (“Count”) and unique CAZyme families (“Families”) are provided for each class, as are the fraction of the total CAZyme count represented by each class (“Fraction of total”), the percent of MAGs with at least one CAZyme in each class (“Presence in MAGs”), the median count of unique CAZymes from each class per MAG (“Median count per MAG”), and the median count of unique CAZyme families from each class per MAG (“Median families per MAG”). Summary data for degradation CAZymes (GH, PL, and CE) as a group are also provided.

|  | Count | Families | Fraction of total (%) | Exported fraction (%) | Presence in MAGs<sup>a</sup> (%) | Median count per MAG | Median families per MAG |
| --- | --- | --- | --- | --- | --- | --- | --- |
| GH | 1972 | 96 | 42.37 | 47.92 | 98.81 | 20.5 | 12.0 |
| PL | 67 | 18 | 1.44 | 67.16 | 39.29 | 0.0 | 0.0 |
| CE | 253 | 13 | 5.44 | 21.74 | 91.67 | 3.0 | 3.0 |
| GT | 1920 | 27 | 41.25 | 3.28 | 100.00 | 23.0 | 10.0 |
| CBM | 386 | 23 | 8.29 | 54.66 | 75.00 | 3.0 | 2.0 |
| AA | 56 | 4 | 1.20 | 7.14 | 33.33 | 0.0 | 0.0 |
| **All** | **4654** | **181** | **100** | **28.4** | **100** | **52.5** | **28.5** |
| Deg.<sup>b</sup> | 2292 | 127 | 49.25 | 45.59 | 100.00 | 24.0 | 15.0 |

<sup>a</sup> The percent of MAGs containing at least one gene from the relevant group.
<sup>b</sup> deg. = GH, PL, and CE together. Collectively called “degradation CAZymes”

All analyzed MAGs had at least one gene associated with carbohydrate degradation. GH, which cleave glycosidic bonds, were the most abundant CAZymes and present in all but one MAG. PL, which break the glycosidic linkages in anionic polysaccharides, were less numerous and present in only about 40% of MAGs (Table 1). MAGs with at least one PL tended to have more degradation CAZymes (i.e., GH, PL, and CE) than those without a PL: 75.7% of MAGs with at least one PL had above-median counts of degradation CAZymes. CE, which catalyze the removal of ester-linked groups such as acetic and carboxylic acids from carbohydrates, were far less numerous than GH but nearly as ubiquitous – they were identified in over 90% of MAGs (Table 1). Over half of all degradation CAZymes contained signal peptides tagging them for export into or out of the cell membrane (Table 1). PL contained signal peptides most frequently (67.2 %), suggesting that PL in this dataset could have roles in polysaccharide degradation or other functions involving extracellular processing of polysaccharides, glycoproteins, and other carbohydrates, such as defense and signaling.

GT were ubiquitous throughout the dataset (Table 1). All MAGs had at least 3 GT. Genes from the three most common GT families, GT2, GT4, and GT51, together accounted for 63.2% of all GT. According to the CAZy database, GT51 (223 genes, 11.6% of GT) is specific for murein (peptidoglycan) transferase activity and is implicated in building the peptidoglycan cell wall of bacteria. GT2 and GT4, in contrast, have a wide range of activities including but not limited to synthesis of sugar-phosphates and oligo- and polysaccharides (22). The 3 Cyanobacteria MAGs had a median of 65 GT genes from a median of 13 distinct families. This was more than double the median count of GTs in non-Cyanobacteria MAGs (22 GT genes from 10 distinct families). Cyanobacteria MAGs each had between 40 and 80 GTs, compared to 2-40 GTs in non-Cyanobacteria MAGs. The elevated numbers of unique GT genes in the Cyanobacteria MAGs suggest that these cyanobacterial GTs could have a greater functional variety within their multifunctional families and/or greater functional redundancy than GTs in other organisms. Both interpretations are consistent with the cyanobacterial production of copious and complex EPS carbohydrates (18, 19).

Owing to the large size of polysaccharides, the first steps of polysaccharide degradation must take place extracellularly or via polysaccharide uptake systems, such as the TonB-dependent system in Bacteroidetes (23–27). Additionally, many CAZyme families are highly specific to only certain glycosidic bonds and substrates. For example, degrading complex carbohydrates such as pectin and fucoidan can require tens or even hundreds of distinct CAZymes (28–30) and special microcompartments for sugar processing such as those of Planctomycetes and Verrucomicrobia (31, 32). Given the complexity and compositional diversity of cyanobacterial EPS, complete EPS degradation in pustular mats should also be expected to require large numbers of functionally varied CAZymes. We hypothesized that MAGs representing putative primary degraders — organisms that initiate extracellular degradation of complex polysaccharide substrates — would be enriched in exported degradation CAZymes relative to other MAGs. Conversely, MAGs representing “downstream” putative EPS degraders that metabolize oligosaccharides released by the degradation of EPS would instead contain large suites of intracellular degradation CAZymes. Furthermore, MAGs representing organisms that specialize in the degradation of a small subset of components of EPS would be expected to have smaller suites of less diverse CAZymes than putative EPS degraders of either group.

We searched for taxonomic trends in the abundance and export tendencies (see Methods) of degradation CAZymes in individual MAGs (Fig. 2) that could identify MAGs as representing organisms as putative primary or downstream polysaccharide degraders. Export tendency was calculated as the signed distance from a MAG to the line with the slope 1 and intercept 0 line normalized to the magnitude of the greatest distance (Methods). The dataset-wide mean export tendency of -0.047 with a standard deviation of 0.30 (p=0.16, N=84 for one-sample t-test with µ= 0) indicated a lack of a strong preference for exported or intracellular CAZymes in the metagenome as a whole.

**Figure 2.**
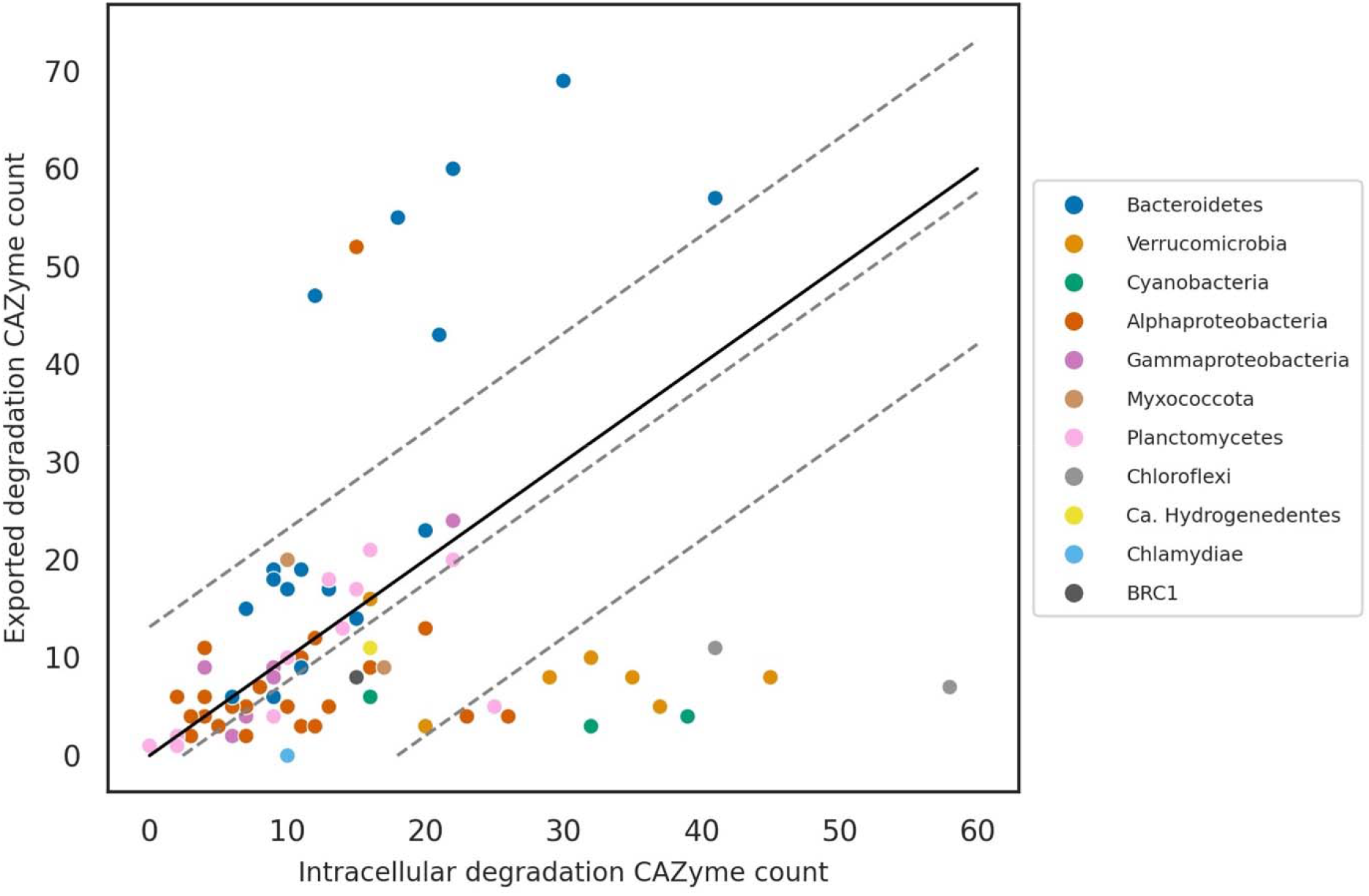
Localization of degradation CAZymes in 84 Shark Bay MAGs. Degradation CAZymes include GH, PL, and CE. Lines: localization preference equal to 0 (solid black), mean localization preference of -0.047 (center gray dashed), localization preference 1 standard deviation above (top gray dashed) and below (bottom gray dashed) the mean. Three distinct clusters of MAGs can be visually distinguished based on this metric.

Figure 2 shows three distinct groups of MAGs with respect to counts and export tendencies of degradation CAZymes. We tentatively assigned them to putative primary EPS degraders, putative downstream EPS degraders, and other organisms. One group, consisting of 6 Bacteroidetes MAGs (2 Saprospirales, 2 Cyclobacteriaceae, and 2 unidentified Cytophagia) and 1 Alphaproteobacteria (Parvularculaceae) MAG had high counts of degradation CAZymes, degradation CAZyme families (59-99 per MAG) and high positive export tendencies (Fig. 2). The second group consisted of 14 MAGs with moderate to high counts of degradation CAZymes and very low export tendencies (i.e., more intracellular than exported degradation CAZymes). This group consisted of 6 Verrucomicrobia (2 Puniceicoccaceae, 2 Chthoniobacterales, 1 Opitutales, and 1 Methylacidiphilales), 2 Chloroflexi (Anaerolineae), 2 Cyanobacteria (both unclassified), 3 Alphaproteobacteria (2 unclassified and 1 Rhodobacteraceae), and 1 Planctomycete (Phycisphaeraceae). All other MAGs fell into a third group with roughly even counts of exported and intracellular degradation CAZymes and mostly below-median total CAZyme counts. Identification of CAZymes tagged for export additionally identified some groups that may specialize in the degradation of specific anionic polysaccharides.

The three groupings reflected differences in phylum-level distributions of the counts and variety of all and exported degradation CAZymes (Fig. 3). Bacteroidetes generally had high counts of degradation CAZymes compared to other groups. Chloroflexi, Verrucomicrobia, and to a lesser extent Cyanobacteria and some Planctomycetes, had abundant degradation CAZymes, but very few of those CAZymes were exported. We observed similar results when considering the number of unique CAZyme families represented in a MAG instead of counts of individual CAZymes. The sole Chlamydiae MAG had *no* predicted exported degradation CAZymes, but unlike Verrucomicrobia and Anaerolineae MAGs, it was also generally very CAZyme-poor, with only 10 CAZymes in total.

**Figure 3.**
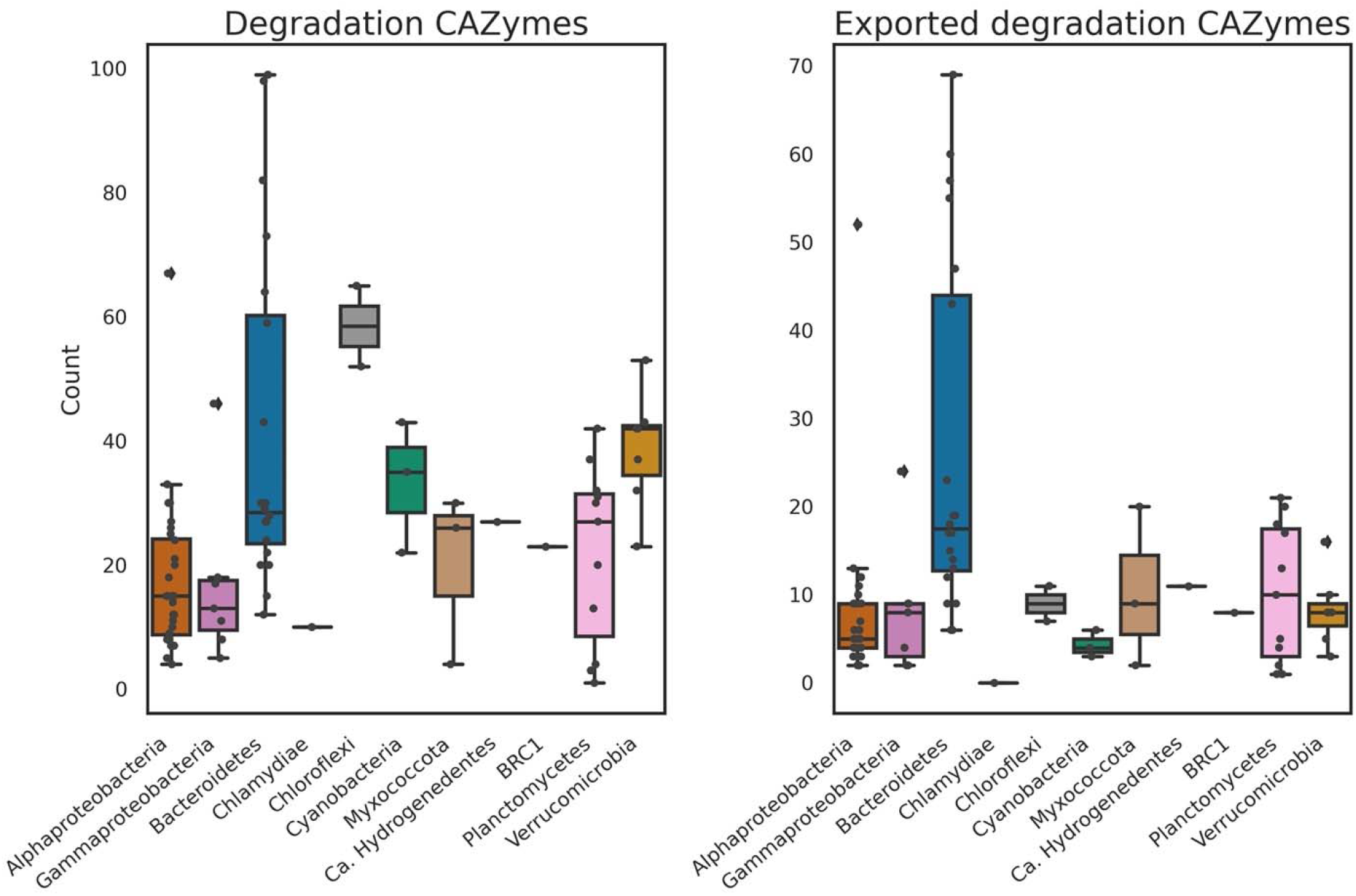
Phylum-level distributions of degradation CAZyme counts in MAGs. Left panel: Total counts. Bacteroidetes (blue) have a high number of degradation CAZymes. Other CAZyme-rich phyla include Verrucomicrobia (tigers eye), Chloroflexi (gray), and Cyanobacteria (green). Right panel: Exported counts. Bacteroidetes have abundant exported degradation CAZymes. Two outliers in Alphaproteobacteria and Gammaproteobacteria have high total and exported CAZyme counts.

Some exceptions to phylum-level trends suggested the presence of microbes with carbohydrate utilization strategies distinct from those typical for their phyla. The counts of degradation CAZymes and the export tendency score in Parvularculaceae MAG 22 were comparable to those of the 5 most CAZyme-rich Bacteroidetes MAGs. This MAG was an outlier among Alphaproteobacteria, which generally had below-median degradation CAZyme counts and export tendencies near 0. Saccharospirillum MAG 65 was similarly set apart from other Gammaproteobacteria by the 8^th^ highest number of exported degradation CAZymes among all MAGs.

GTs are generally associated with carbohydrate synthesis rather than degradation and their localization did not exhibit similar groupings. Only 3.3% of GTs identified in the metagenome contained signal peptides (Table 1) and did not seem to be concentrated in any taxonomic group. The few MAGs with exported GTs were scattered across most phyla represented in the metagenome, with at least one exported GT in all three Cyanobacteria MAGs. Those MAGs that did contain exported GTs most often did not have more than one exported GT from a single family, indicating potential functional promiscuity of enzymes encoded by these gene families. This is consistent with the association of many GT families with numerous activities according to the CAZy database (22).

To understand the minimum potential diversity of polysaccharide and oligosaccharide substrates for CAZymes in pustular mats, we considered the predicted substrates of GH and PL and looked for taxonomic trends in predicted substrate utilization potential (Figs. 4 and 5). CAZymes from both groups cleave glycosidic bonds and sequence-based families of these enzymes often have very specific substrates (Table 2). We excluded 7 polyspecific GH families (343 GH genes and 17% of total GH) with predicted activity on more than 4 distinct glycosidic bonds (e.g., alpha 1-3 glucosidic bond) or more than 9 substrates (e.g., alpha glucans) from the analyses of predicted substrates to avoid diluting the more reliable signals of more specific CAZymes. Figures 4 and 5 show these polyspecific families as their own “substrate” categories. Overall, CAZymes in Shark Bay pustular mats are likely to have an even broader diversity of potential substrates than predicted here.

**Table 2.** Potential substrates of GH and PL identified in a Shark Bay pustular mat metagenome. Counts of GHs in each substrate category are equal to the number of GH genes identified in families associated with that substrate category. Categories were assigned to gene families using a semi-automatic process based on CAZydb annotations and EC numbers (see Methods). Single families may be associated with multiple substrates, but polyspecific families (see Methods) were excluded from these counts.

| GH substrate | GH count | families |
| --- | --- | --- |
| alpha-glucan | 432 | GH13, GH133, GH14, GH15, GH57, GH63, GH77, GH97 |
| beta-glucan | 427 | GH10, GH128, GH13, GH133, GH144, GH148, GH149, GH158, GH16, GH17, GH26, GH30, GH48, GH51, GH6, GH8, GH81, GH9, GH94 |
| peptidoglycan | 318 | GH102, GH103, GH104, GH108, GH18, GH19, GH23, GH24, GH25, GH73 |
| chitin | 243 | GH108, GH18, GH19, GH23, GH24, GH25, GH48, GH73, GH94 |
| xylan | 140 | GH10, GH115, GH116, GH120, GH141, GH26, GH30, GH39, GH43, GH51, GH62, GH67, GH8 |
| mannan | 121 | GH113, GH130, GH26, GH36, GH57, GH97 |
| galacturonan | 84 | GH105, GH106, GH138, GH28, GH78, PL1, PL10, PL11, PL9 |
| arabinogalactan | 84 | GH30, GH39, GH43, GH51, GH53, GH62 |
| alpha-galactan | 81 | GH36, GH57, GH97 |
| arabinan | 76 | GH39, GH43, GH51, GH62 |
| glycosaminoglycan<sup>a</sup> | 70 | GH16, GH166, GH39, GH79, GH84, PL12, PL15, PL29, PL33, PL35, PL6, PL8 |
| beta-galactan | 55 | GH16, GH30, GH43 |
| algal polysaccharides<sup>b</sup> | 37 | GH117, GH16, GH168, GH50, GH86, PL14, PL15, PL17, PL31, PL40, PL5, PL6, PL7 |

<sup>a</sup> chromatin sulfate, dermatan sulfate, heparin/heparan sulfate
<sup>b</sup> fucan, carrageenan, beta-(1,4)-glucuronic acids, algin (not necessarily sulfated), and ulvan

**Figure 4.**
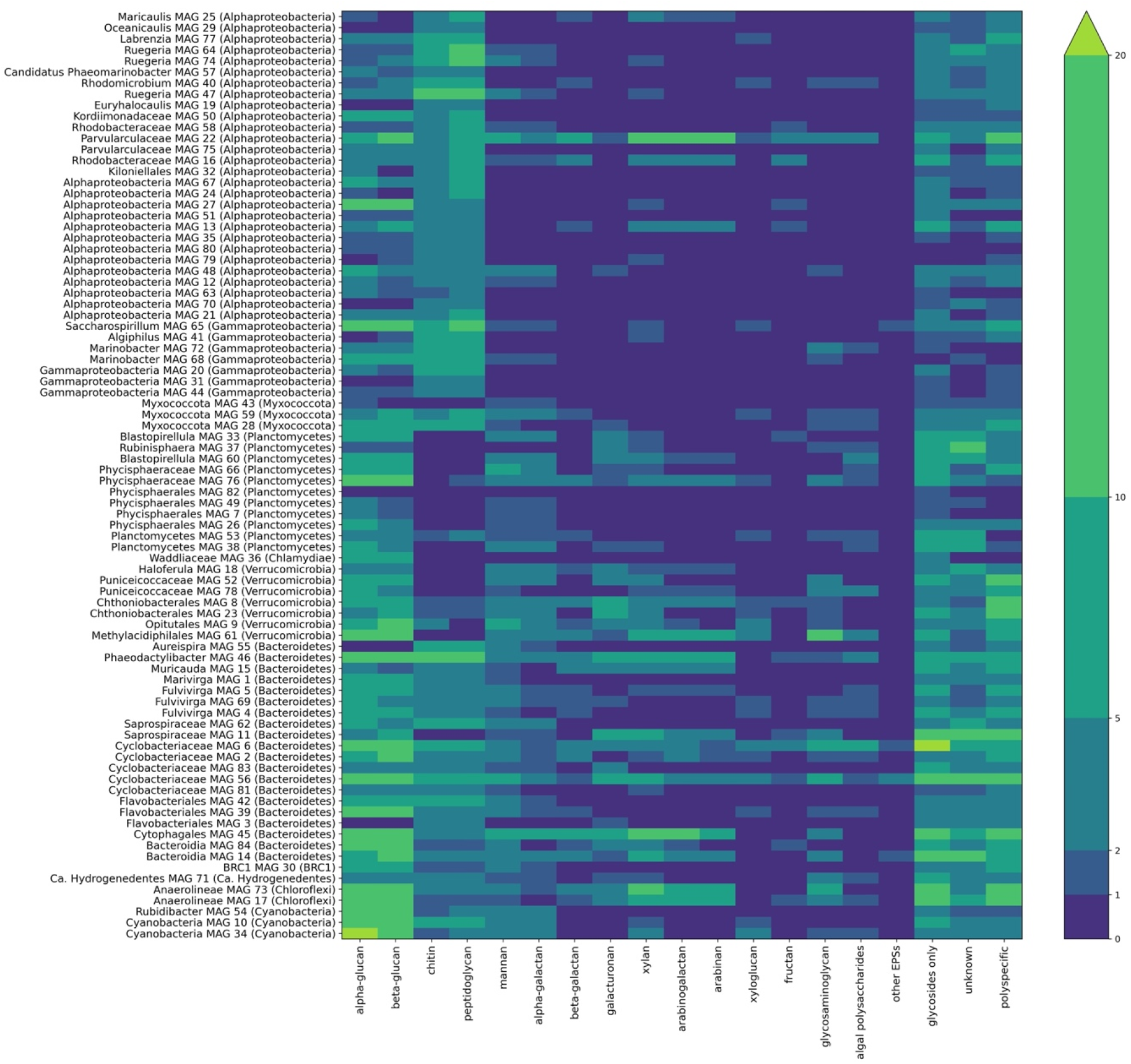
Distribution of potential polysaccharide substrates of GH and PL identified in Shark Bay MAGs. Colors reflect the number of genes in a given MAG with predicted activities on respective substrates. Genes can be counted more than once if they have more than one potential substrate. Genes from polyspecific families were counted only in the column labeled “polyspecific” (see Methods). The abundance of GH with predicted activity only on glycosides (i.e., not specifically on polysaccharides or oligosaccharides) are shown in the column labeled “glycosides only.” GH and PL with unknown activities are shown in the column labeled “unknown.” Algal polysaccharides include algin, carrageenan, ulvan, fucan, and beta-(1,4)-galacturonan. Glycosaminoglycans (GAGs) include chondroitin sulfate, dermatan sulfate, heparin/heparan sulfate, and unidentified GAGs. “Other EPSs” include the known bacterial exopolymers xanthan, gellan, and polygalactosamine.

**Figure 5.**
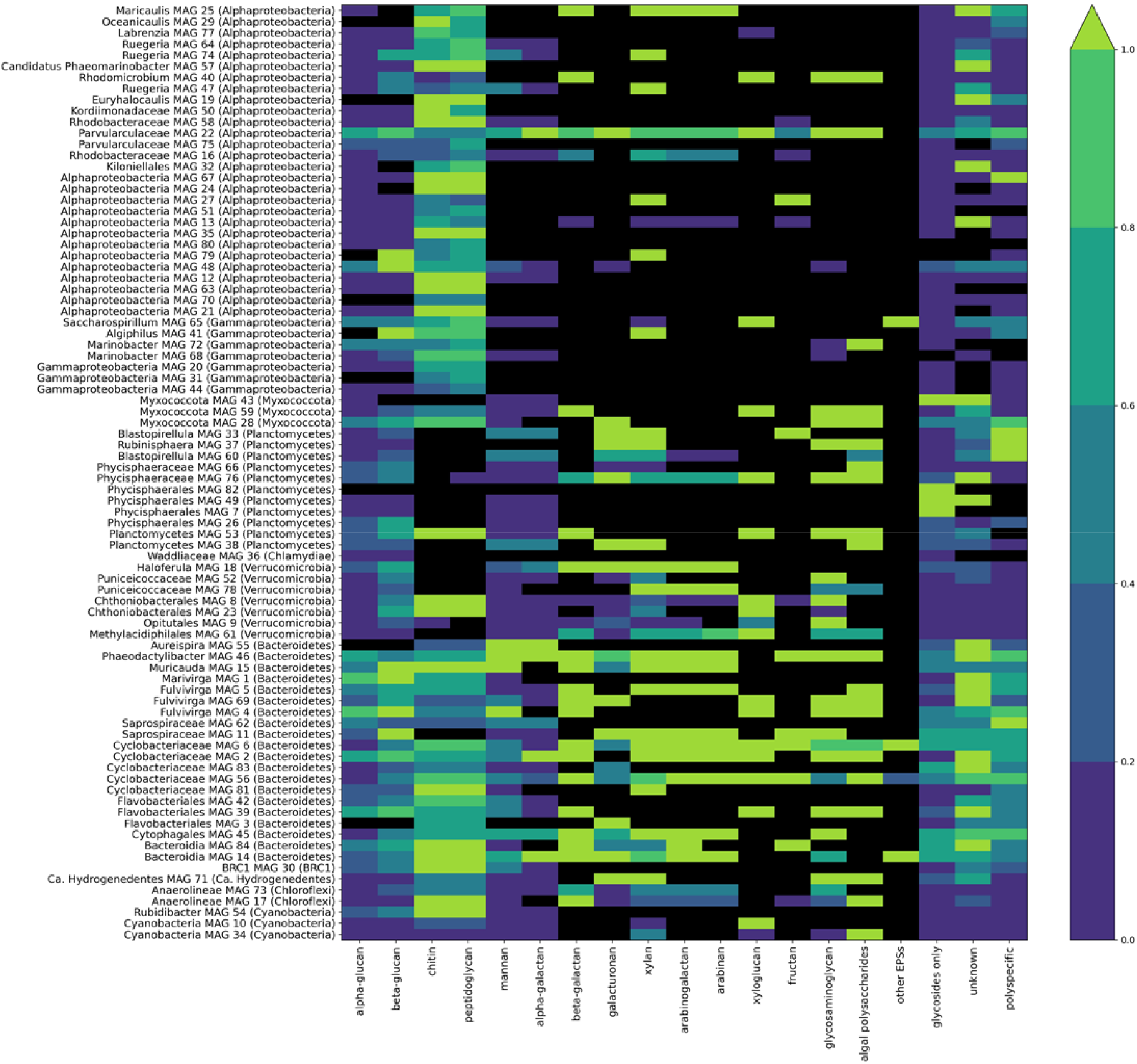
Proportion of exported GH and PL by predicted substrates. Substrate categories and polyspecific families are treated as in Figure 4. For each MAG (rows) and predicted GH/PL substrate (columns), the colored areas represent the proportion of associated GH and PL genes that is predicted to be exported. The color scale is shown on the right. Black areas denote the lack of detected CAZymes (exported or intracellular) associated with a given substrate from individual MAGs.

Nearly all microbial taxa represented by the 84 MAGs from pustular mats have the potential to utilize carbohydrates that contain D-glucose. These taxa also possess genes for modifying or breaking down bacterial cell walls (which could include their own) and potentially chitin (Fig. 4; Table 2). CAZyme families with predicted activity on peptidoglycan often contain genes with predicted activity on chitin and vice-versa, complicating confident discrimination between the two predicted substrates. Predicted peptidoglycanases and chitinases were notably absent from most MAGs in the Planctomycetes, Verrucomicrobia, and Chlamydiae (PVC) superphylum that otherwise had moderately high CAZyme counts. This absence may reflect some aspect of the unusual and controversial nature of peptidoglycan cell walls in PVC superphylum (33, 34, 34– 37).

CAZymes targeting other carbohydrates tended to be less numerous and more taxonomically restricted and appeared mainly in groups with high CAZyme counts and family diversity. Of particular interest, given the suggested roles in mineral precipitation (9, 18, 38–40). The many anionic polysaccharide substrates predicted for CAZymes in our dataset. These included sulfated polysaccharides such as sulfated glycosaminoglycans (GAGs) and algal polysaccharides as well as polysaccharides containing uronic acids, such as galacturonans, the algal polysaccharides algin and beta-(1,4) glucuronan, and the bacterial EPSs xanthan and gellan. Most Bacteroidetes, Verrucomicrobia, Chloroflexi, and Planctomycetes MAGs had at least one GH or PL with predicted anionic substrates. Only 5 of the 35 Proteobacteria MAGs contained a GH or PL with predicted activity on a sulfated GAG, galacturonan, or algal polysaccharide. Additionally, about 10% of GH and PL (259 total) were unidentified, suggesting an even greater variety of possible carbohydrate degradation functions in Shark Bay mats than reported here.

The tendency of GH and PL with certain predicted substrates to be exported (Fig. 5) suggested the presence of these substrates in the extracellular matrix. For example, CAZymes with the predicted roles in depolymerizing galacturonans were mostly exported in Bacteroidetes (Fig. 5). Most CAZymes targeting beta-galactans contained export sequences, except for those in Verrucomicrobia MAGs. Most MAGs had at least one GH or PL with predicted activity on glucans, peptidoglycan, chitin, mannans, and alpha-galactans, but GH and PL with other, “rarer” predicted substrates were less widespread and typically contained sequences targeting these enzymes for export (Fig. 5). These GH and PL were mostly exported and most abundant in Bacteroidetes, Verrucomicrobia, and Chloroflexi MAGs, and also present in about half of the Planctomycetes MAGs. Parvularculaceae MAG 22 (Alphaproteobacteria) had many GH and PL acting on these substrates (Fig. 5), distinguishing it from most other Alphaproteobacteria MAGs. Rhodomicrobium MAG 40 also contained excreted CAZymes predicted to degrade algal polysaccharides, but contained overall fewer CAZymes compared to MAG 22 and to other CAZyme-rich MAGs (Fig. 2). Additionally, the one Hydrogenedentes MAG had slightly above-median counts of GT (24) and degradation CAZymes (27), a moderately above-median number of distinct degradation CAZyme families (20) and more degradation CAZymes that were not tagged for export, including enzymes with predicted activity on sulfated substrates (Figs. 4 and 5).

CE may also contribute to polysaccharide degradation in Shark Bay pustular mats. Because the removal of ester-linked groups from carbohydrates by these enzymes is distinct from the activities of GH and PL, and because CE families often lack the detailed substrate-level annotation, we considered them separately from PL and GH. All of the 253 CE identified had predicted deacetylase activity. Approximately one third of the identified CE belonged to just four families: CE11, CE4, CE14, and CE9, all with predicted N-acetylglucosamine deacetylase activities. CEs belonging to other families were most common in Bacteroidetes, Planctomycetes, Chloroflexi, and Verrucomicrobia, echoing the greater diversity of “rare” GH and PL in these groups. Only one carboxyl esterase family was identified, CE1, with additional acetylesterase activity. 30 of the 84 MAGs had at least one CE containing a signal peptide tagging it for export. This included 9 of 11 Planctomycetes MAGs, 5 of 7 Verrucomicrobia MAGs, 7 of 20 Bacteroidetes MAGs, 1 of 3 Cyanobacteria MAGs, 4 of 28 Alphaproteobacteria MAGs including Parvularculaceae MAG 22, and 1 of 8 Gammaproteobacteria MAGs (Saccharospirillum MAG 65). These analyses predict acetylation of polysaccharides in pustular mats and support the roles of Bacteroidetes, Planctomycetes, Chloroflexi (Anaerolineae) and Verrucomicrobia, as well as the CAZyme-rich MAG 22 and MAG 65 in the degradation of acetylated EPSs.

### Microbial communities enriched on different polysaccharides

The distribution of the genes encoding for diverse CAZymes among the MAGs (Figs. 4, 5) predicted that different microbial groups might grow and become enriched on different polysaccharides. To test this, we inoculated liquid media amended with six different polysaccharides with pustule material, grew the resulting cultures in the dark for two weeks and analyzed the composition of enrichments by sequencing and analyzing 16S rRNA gene amplicons. The compositions of microbial communities present in the dark polysaccharide enrichment cultures were compared to those of the inoculum material (“slurry” of crushed pustule material suspended in liquid media) grown in illuminated polysaccharide-free media and to raw pustule material that had been stored in a dark refrigerator at 4 °C (Fig. 6). Cultures with visible turbidity yielded an average of 377,306 reads, compared to an average of 49,117 reads in the cultures that did neither became turbid nor yielded detectable DNA upon extraction (i.e., that did not grow). We identified 3602 amplicon-sequence variants (ASVs) in total. The pustule community was by far the most diverse, with 3180 ASVs and the pectin and chitin enrichments that did not grow had the lowest diversity, with 47 and 58 ASVs present in these enrichments, respectively. A large fraction of slurry ASVs (69 of 205) were found in enrichments that grew (i.e., agar, cellulose, chondroitin sulfate, laminarin, and xylan).

**Figure 6.**
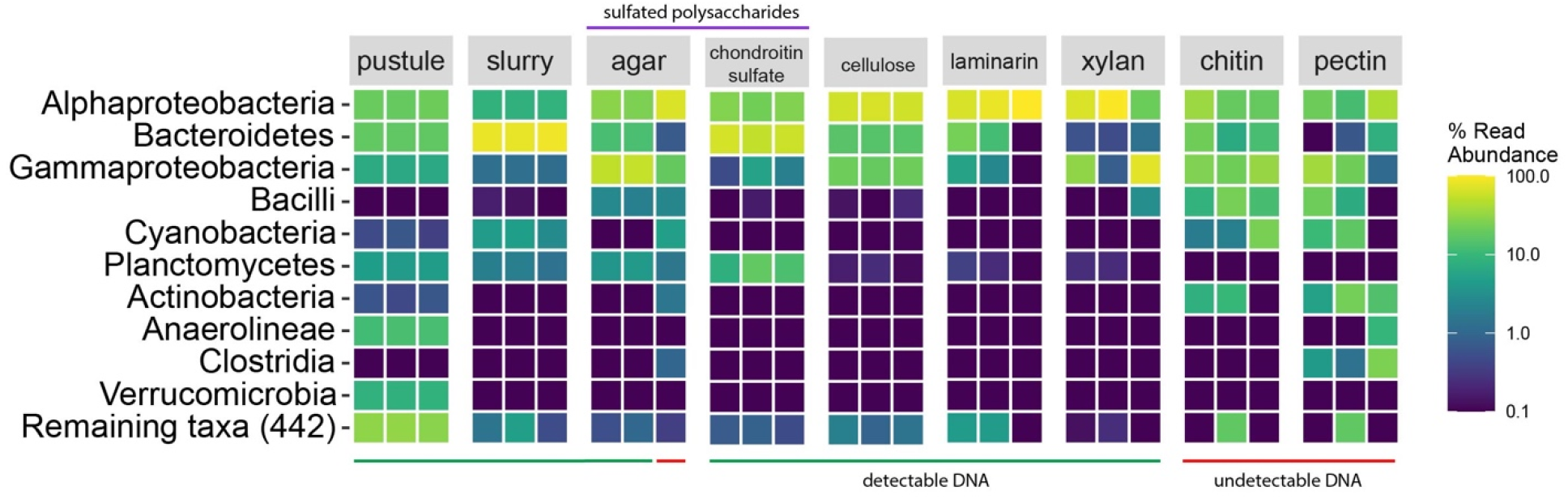
Relative abundances of ASVs in polysaccharide enrichment cultures derived from a Shark Bay pustular mat. Pustule is the original frozen mat material thawed before establishing the slurry and then stored in the dark at 4 °C for the duration of the experiment. Slurry is the inoculum, crushed thawed pustule suspended in hypersaline BG-11 media and grown in the presence of light. Columns labeled agar, cellulose, chondroitin sulfate, laminarin, xylan, chitin, and pectin show data from enrichment cultures inoculated with crushed pustules and incubated in the dark in hypersaline BG-11 media amended with the corresponding polysaccharide (e.g., agar, cellulose). Each square corresponds to one biological replicate. Color bar shows percentages of read abundances. The red line indicates cultures for which DNA extraction yields fell below detection limits (i.e., that did not grow). The purple line indicates sulfated substrates.

Microbial communities from pustular mats were able to grow on diverse polysaccharides that were predicted as potential substrates based on MAG analyses: cellulose, laminarin, agar, chondroitin sulfate, and xylan. The raw pustule and the light-grown slurry contained abundant cyanobacterial ASVs, but none of the polysaccharide enrichments whose turbidity increased in the dark did. The community compositions of enrichments on cellulose, chondroitin sulfate, laminarin, and xylan, as well as two of the three agar enrichments, were distinct from the communities in the original pustules and from the illuminated slurry. The communities enriched on these polysaccharides were dominated by Alphaproteobacteria, Gammaproteobacteria and Bacteroidetes. Sulfated polysaccharides (i.e., agar and chondroitin sulfate) also enriched for Planctomycetes. The slurry and all enrichment cultures that yielded measurable DNA concentrations upon extraction grouped apart from the low-DNA cultures and pustule on a PCoA ordination plot of Bray-Curtis dissimilarity (41) (Fig. 6). Most of the enrichments that exhibited visible turbidity after the growth period grouped together, with chondroitin sulfate and the slurry forming distinct clusters (Fig. 7). Chondroitin sulfate grouped closest to the slurry, which suggests closer compositional and structural similarities of GAG to EPS polysaccharides relative to other tested polysaccharides.

**Figure 7.**
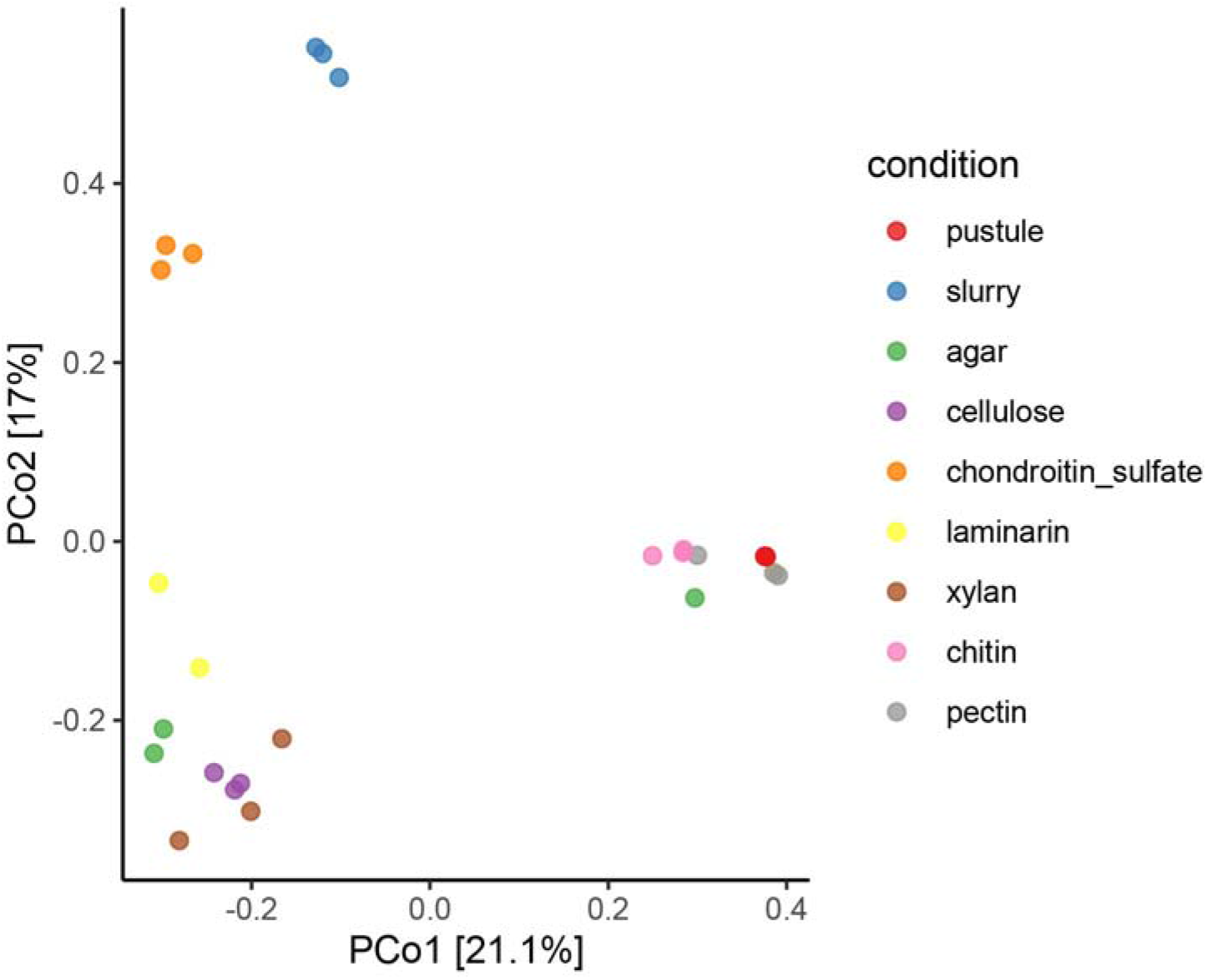
Principal Coordinates Analysis (PCoA) based on the Bray-Curtis distance measure of 3602 ASVs and 27 samples ASVs derived from samples and polysaccharide enrichment cultures inoculated by dispersed pustular mat from Shark Bay. Colors indicate the source of the ASVs (see Figure 6 for the descriptions of conditions). The relative contribution (eigenvalue) of each axis to the total inertia in the data is indicated in percent at the axis titles.

Alphaproteobacteria had a greater relative abundance in most of the polysaccharide enrichments compared to the pustule and slurry. Alphaproteobacterial ASVs of the genera *Pelagibacterium, Parvularcula*, and *Nesiotobacter* dominated the Alphaproteobacterial community in enrichments on glucans (i.e., laminarin and cellulose) and xylan. *Pelagibacterium* was particularly dominant in enrichments on laminarin (Fig. 6). The enrichment of *Parvularcula* is notable given that the anomalously CAZyme-rich Alphaproteobacterial MAG 22 we recovered from the Shark Bay pustule belonged to Parvularculaceae. While present, none of these genera were detected as abundant members of the pustule or slurry communities and were also less dominant in the communities enriched on sulfated polysaccharides.

Gammaproteobacterial ASVs mostly belonged to the genera *Marinobacter* across all enrichments and the slurry and were more abundant overall in the polysaccharide enrichments compared to the illuminated slurry. Two *Marinobacter* MAGs were also assembled from the pustular mats. Overall, taxa that were abundant in the pustule were not present at high abundances in the dark amended enrichments or in the slurry. Groups of Alphaproteobacteria and Gammaproteobacteria that were abundant in the enrichments on single polysaccharides were likely outcompeted by other organisms, namely Bacteroidetes, when growing on complex substrates such as cyanobacterial EPSs in pustules and the slurry.

Bacteroidetes dominated the community in the illuminated slurry, were less abundant but still a major constituent of the community in most of the enrichment cultures that grew (>10 %), and were, on average, only 0.81% of the community in the xylan enrichments. Bacteroidetes ASVs in both the slurry and enrichments were dominated by *Marivirga* (Cytophagales) and *Muricauda* (Flavobacteriales). These organisms belong to the same families as many Bacteroidetes MAGs, including one unidentified Cytophagales, unidentified Cyclobacteriaceae (Cytophagales) and Flavobacteriaceae. Less abundant Bacteroidetes genera varied across the various growth conditions, with *Salegentibacter, Cryomorpha*, and *Imperialibacter* accounting for between 0 and ∼10% of the Bacteroidetes ASVs in the agar, cellulose, chondroitin sulfate, and laminarin enrichments. The slurry community hosted a greater number of distinct Bacteroidetes ASVs than the enrichments on polysaccharides. Among the enrichments, the chondroitin sulfate enrichment contained the highest percentages of Bacteroidetes and Planctomycetes ASVs. The latter ASVs accounted for <1% of the community in enrichments on non-sulfated polysaccharides, increasing to 12.8% of the community in the chondroitin sulfate enrichments and 3.8% of the community in agar enrichments (excluding the low-DNA replicate). Bacteroidetes reached an average relative abundance of 58.8% in the chondroitin sulfate enrichment and 12.5% in the agar enrichments with growth. This observation is consistent with the large number of exported CAZymes with predicted sulfated and anionic substrates in Bacteroidetes and Planctomycetes MAGs.

Neither the chitin nor pectin enrichment cultures exhibited visible growth after two weeks. The concentrations of DNA extracted from these cultures and one of the agar enrichment triplicates were below the detection limit (Fig. 6). Most of these enrichments contained cyanobacterial ASVs despite the dark incubation, indicating that very little growth occurred in these cultures and that their community compositions reflected relict DNA from the pustule material inoculum. PCoA ordination, which grouped the seven low-DNA replicates with pustule material, further supports this inference (Fig. 7).

## Discussion

Delineating specific roles of different microbial groups in EPS degradation is a vital step in interpreting microbialites as products of interactions between microbes and the organic matrix that houses microbial cells in tidal environments. Our results demonstrate expansive suites of CAZymes in the pustular mats from Shark Bay and experimentally substantiate roles for different microbial groups in carbohydrate synthesis and degradation. Anomalously high GT counts and diversity in Cyanobacteria MAGs confirm a role in polysaccharide production for Cyanobacteria. Some major phyla that are abundant and ubiquitous in mats, such as Bacteroidetes, Planctomycetes, and some Proteobacteria, are likely involved in initial polysaccharide degradation, with Bacteroidetes and Planctomycetes being particularly enriched by sulfated and cyanobacterial polysaccharides. These analyses predict the roles for Verrucomicrobia and Anaerolineae in downstream degradation of diverse carbohydrates potentially derived from EPS as well as in the degradation of sulfated EPS and is consistent with the presence of sulfatase genes in these organisms (19, 30). Errors in the binning and assembly of metagenomes can inflate the counts of distinct CAZyme genes within individual MAGs and may therefore have impacted our conclusions. However, the phylum-level trends we observe hold even when considering the diversity of families alone (i.e., presence/absence of specific CAZyme families). In fact, even the highest degradation CAZyme counts in our MAGs are consistent those of known polysaccharide-degrading organisms (30, 42–44, 44–47). Additionally, some common CAZyme families encompass genes with a wide range of known possible activities (e.g., GT4, GH2) (22), so collapsing our analysis to family diversity would have obscured an important potential indicator of diverse functional potential in the MAGs.

The chemical structures of marine cyanobacterial EPSs are mostly unknown, but it is generally understood that these diverse polymers are usually very large (>1 MDa) and complex, commonly incorporating 6-13 different sugars (including uronic acids, amino sugars, and deoxysugars), in contrast to other bacterial EPSs typically contain less than 4 (48). Accordingly, we observed that Cyanobacteria MAGs all possessed a much greater number and variety of GTs than all other MAGs. This is also consistent with previous studies showing that GTs are enriched in the upper layers of Shark Bay mats where Cyanobacteria are most abundant and active (14, 20).

The polysaccharide degradation potential of heterotrophic MAGs also reflects the complexity of EPS carbohydrates: we found that bacterial MAGs with high numbers of degradation CAZymes contained GH and PL with predicted activity on polymers of the above mentioned sugars. Most MAGs, even those without many other CAZymes, possessed at least a few export-tagged genes for processing glucan, suggesting that glucans are a component of the EPS matrix in Shark Bay pustular mats, as they are in the cyanobacterial mats of Elkhorn Slough (49). Beta-glucosidases and alpha-glucosidases also had measurable activities in the hypersaline microbial mats on Eleuthera Island, The Bahamas (10).

Many MAGs, and particularly Bacteroidetes, Planctomycetes, Verrucomicrobia, Chloroflexi, Myxococcota, Hydrogenedentes and Parvularculaceae MAG 22, also had CAZymes predicted to degrade sulfated polysaccharides, compounds recently confirmed in Shark Bay pustular mat EPS (19) and also likely present in the hypersaline mats in Eleuthera Island (10). CEs predicted to be exported into the periplasm or outside the cell were found almost exclusively in Bacteroidetes, Planctomycetes, and Verrucomicrobia MAGs and in a few Alphaproteobacteria and Gammaproteobacteria MAGs with overall high counts of exported degradation CAZymes. This distribution suggests that not only sulfated, but also acetylated polysaccharides could be a component of Shark Bay EPSs.

Trends in the abundance, diversity, and export tendency of CAZymes in the Shark Bay pustular mat MAGs suggested distinct roles in EPS cycling for many ubiquitous phyla in these mats. Microbial communities enriched on some predicted substrates supported these roles. Cyanobacteria had abundant, mostly intracellular degradation CAZymes and grew in our culture experiments only under light. We therefore interpret their large, diverse suites of intracellular degradation CAZymes as being involved in processing and modifying their own carbohydrate stores and EPSs (21, 49–53). Given that Verrucomicrobia and Anaerolineae also had abundant and varied intracellular CAZymes, these organisms may degrade oligosaccharides that are released by organisms that perform the first extracellular steps of EPS polysaccharide degradation. Both groups also possessed some exported degradation CAZymes, particularly for the degradation of sulfated polysaccharides, and therefore might also participate in primary EPS degradation. Future transcriptomic work will be required to definitively determine which organisms express exported proteins. Anaerolineae are core members of cellulolytic bioreactor communities, though their own ability to directly degrade cellulose can be limited, consistent with a potential downstream role in EPS degradation (54, 55). Verrucomicrobia are found in high-carbohydrate systems such as human and animal guts (56, 57), rice paddies (58), organic-rich lakes (59) and marine macro- and microalgae (60–62), and are functionally and numerically important members of bacterioplankton communities that degrade recalcitrant sulfated algal polysaccharides including fucoidan (30, 63). Verrucomicrobia also have specialized sugar-processing microcompartments similar to those of Planctomycetes (31, 32). Some Verrucomicrobia MAGs contained predicted exported xylanases, alpha-glucanases and beta-glucanases, in keeping with previously reported exported laminarase and xylanase in polysaccharide-degrading marine Verrucomicrobia (43). Genes containing Planctomycete-specific cytochrome *c* (PSCyt) domains (pfam07635, pfam07583, and pfam07627) have been reported in Verrucomicrobia MAGs representing likely saccharolytic organisms, where they tended to occur in the genetic neighborhood of sulfatases and carbohydrate modules and were attributed a possible function in intracellular degradation of carbohydrate mono- and oligomers (59). Genes containing PSCyt domains were found in all but one Verrucomicrobia MAG and in all but two Planctomycete MAGs, further supporting roles for these phyla in EPS degradation in Shark Bay pustular mats. Notably, only Planctomycetes were enriched in liquid cultures amended by agar and chondroitin sulfate, suggesting that additional factors control the growth of Verrucomicrobia in natural mats. One such factor could be the presence of oxygen — we allowed air to circulate in the enrichment cultures and did not enrich Anaerolineae, which are obligate anaerobes, either.

The large, varied, and mostly exported CAZyme suites of Bacteroidetes—including CAZymes with predicted activity on sulfated substrates—and their greater relative abundances in the light-grown slurry culture that also enriched for Cyanobacteria and in the dark culture amended by chondroitin sulfate is consistent with a function of Bacteroidetes as primary EPS degraders of various sulfated and other acidic cyanobacterial polysaccharides. We do not find it likely that Bacteroidetes were specifically enriched under light because of proteorhodopsins (PRs) (64):14 of Bacteroidetes MAGs contained BLAST hits (E < 1e-5) to a PR protein sequence from the marine Bacteroidetes *Polaribacter sp. MED152* (GenBank: EAQ40925.1), as did *Marinobacter* MAG 68 (Gammaproteobacteria) and 2 Alphaproteobacteria (MAGs 27 and 48). Instead, the functional potential and observed enrichment patterns of Bacteroidetes echo the known role of Bacteroidetes as degraders of complex polysaccharides in other systems such as the human gut (26, 29, 65–67). Compared to specific Alphaproteobacteria and Gammaproteobacteria that grew in most enrichment cultures, Bacteroidetes may be better equipped to degrade structurally and compositionally complex cyanobacterial EPSs but were likely outcompeted by Proteobacteria in cultures amended by single, non-sulfated polysaccharides.

Previous work suggests a tight link between cyanobacterial EPS production and degradation by Bacteroidetes. Earlier studies identified co-enrichments of Bacteroidetes and Cyanobacteria in the same regions of pustular mats (14) and suggested that Bacteroidetes and Alphaproteobacteria most actively produce mRNA transcripts during the day, when Cyanobacteria photosynthesize (68). The extensive functional potential of Bacteroidetes to perform the first steps in degrading fresh cyanobacterial EPS may explain these trends and link Bacteroidetes to other important metabolic guilds in mats. For example, the flow of carbon from Bacteroidetes to Deltaproteobacteria that consume sulfate and low molecular weight carbon could explain why Bacteroidetes and Deltaproteobacteria were recently found to be the two most active taxa in EPS-rich “cobbles” derived from intact mats disturbed by cyclones (69).

Our results also identify Planctomycetes, Verrucomicrobia, Anaerolineae, Myxococcota and some Alphaproteobacteria, Gammaproteobacteria and Hydrogenedentes as additional potential primary degraders of sulfated EPS. Planctomycetes were enriched on sulfated polysaccharides in our experiments and are an especially promising target for future studies of sulfated EPS cycling in mats. These trends are consistent with the known role of some marine Planctomycetes in the uptake and degradation of sulfated algal and animal polysaccharides (27, 70, 71) and the discovery of specialized sugar-processing microcompartments in Planctomycetes (31). Planctomycetes were enriched in microbial communities grown on chondroitin sulfate and these communities clustered closest to the illuminated slurry in our PCoA. These observations support both the affinity of Planctomycetes for sulfated EPS and compositional similarities between the cyanobacterial EPS (18, 48) and chondroitin sulfate. Chondroitin sulfate, a sulfated animal polysaccharide that consists of an alternating chain of N-acetylglucosamine and glucuronic acid moieties decorated with sulfate groups, is not likely to be an abundant substrate in the mats because high salinity limits the animal community in the hypersaline Hamelin Pool (72). However, cyanobacterial EPS in pustular mats is sulfated (73) and uronic acids have been reported to occur 90% of cyanobacterial EPSs (74), so they can be expected in pustular mats as well. Given this, several genes with predicted activity solely on animal glycosaminoglycans might be reasonably interpreted as acting on structurally similar sulfated polysaccharides produced by mat microbes.

Our comprehensive assessment of CAZymes and their potential substrates in the metagenome of a carbonate-precipitating marine mat cannot illuminate the relative “importance” of microbial groups to gene expression or substrate turnover. Additionally, there are comparably few characterized CAZymes acting on marine-specific substrates such as algal and cyanobacterial polysaccharides that can be used to inform sequence-based functional annotation (52, 75, 76), and our results indicate that some genes could be acting on substrates that are yet to be linked to their CAZyme families. Further, many GH and PL had no known predicted substrate at all. Additionally, exoenzymes secreted by primary degraders may remain bound to the EPS matrix, where they could potentially be utilized by organisms that themselves do *not* excrete export enzymes. This would extend the polysaccharide utilization functions of mat microbes beyond the limits of their own genomes. In this case, putative primary degraders would act as “libraries” of carbohydrate-processing functions that can be “borrowed” by the community via the enzyme-laden matrix. Future transcriptomic studies will be needed to determine the activities and genes of specific organisms and groups of organisms that participate in EPS degradation.

The metagenomic and experimental results presented here generate and support new hypotheses about the ecology of EPS cycling in pustular mats and serves as blueprint for future studied aimed at quantifying and verifying the specific contributions of microbes to EPS degradation. Specifically, integrated models of microbial community structure, function and biomineralization in these ecosystems need to consider EPS-centered interactions and the activities of specific microbial groups and enzymes that modify acidic polysaccharides.

## Materials and Methods

### Metagenomic investigation of EPS-cycling potential

We assessed the genetic potential for cycling of EPS polysaccharides in a set of 84 medium-to-high quality metagenome-assembled genomes (MAGs) derived from a Shark Bay pustular mat.

#### Carbohydrate-active enzyme annotation and substrate assignment

Carbohydrate-active enzymes (CAZymes) were annotated in detail, mapped to predicted substrates and analyzed for the signal peptide secretion tags to establish quantitative patterns in CAZyme distribution, substrate preferences, and excretion patterns among the 84 MAGs. Glycoside hydrolase (GH), polysaccharide lyase (PL), carbohydrate esterase (CE), and glycosyl transferase (GT) genes were annotated according to the classification scheme of the CAZy database (22) using a local installation of the dbCAN2 web server annotation pipeline (77). The pipeline integrates three tools to identify CAZymes: HMMER3 (78) to search against the dbCAN CAZyme HMM database (79), DIAMOND (80) to find Blast hits to the CAZy database, and a Hotpep (81) search for sort conserved CAZyme motifs. All searches were performed with default parameters using the run_dbCAN 2.0.6 Python package (https://github.com/linnabrown/run_dbcan). CAZymes were assigned to families based on the agreement of at least 2 of the 3 search tools (77).

GH and PL families were linked to predicted functions and substrates by assessing the categories of substrates (e.g., alpha-glucan) and bonds (e.g., beta-(1,3) glucose) targeted by each family. To start, specific substrates (e.g., chitin, beta-(1,4)-glucan) and activities (e.g., endohydrolysis of alpha-(1,4) bonds between glucose monomers) were manually assigned to EC numbers as provided by the CAZyme database. Then, these manual associations were used to populate CAZyme families with substrates and activities based on their EC numbers (Supplemental Data S1, Supplemental Data S3). Substrate and activity assignments were then refined by cross-referencing the automatically-generated assignments against the CAZydb activity description strings. Sets of categories for substrates (i.e., alpha-glucanase) and bond activity (alpha-glucosidase) were defined to more succinctly summarize the potential activities observed in the dataset (Supplemental Data S2, Supplemental Data S3). GH and PL were considered polyspecific if they had predicted activity on more than 4 different specific glycosidic bonds or 9 different substrates.

#### Signal peptides and export tendency metric

All annotated CAZymes were submitted to the SignalP version 5.0 web server for the prediction of signal peptides that tag gene products for transport into or out of the cell membrane (82). CAZymes predicted to contain a signal peptide are termed “exported” throughout this paper. Because some MAGs had 0 values for either exported or intracellular CAZyme counts, using a simple ratio of exported to intracellular CAZymes would sometimes result in zero or infinite values, confounding dataset-wide analysis. To avoid this complication, we defined *export tendency* as the signed distance from the line with slope 1 and intercept 0 on the plot of exported vs. intracellular CAZyme counts (Fig. 2). The values were normalized such that the largest observed magnitude was equal to 1. Large positive export tendency indicated more exported than intracellular CAZymes and large negative numbers indicated the opposite. Values near zero reflected a relatively even partition between exported and intracellular CAZymes.

#### Metagenomic data analysis

The outputs of run_dbCAN and SignalP were processed using a combination of Python scripts and Jupyter Notebooks. A directory containing the code for data processing and analysis of the dbCAN and SignalP output can be found at https://github.com/emcuttsy/shark-bay-cazymes.

### Polysaccharide enrichment experiments

Material derived from Shark Bay pustular mats was incubated with a range of polysaccharide substrates and the resulting shifts in community composition quantified by 16s rRNA gene amplicon sequencing. Samples of Shark Bay pustular mats had been previously collected on Carbla Beach in 2017 under license Regulation 17 License #08-000373-1 (19). These samples had been stored in the dark at -20ºC since transport to the lab. Some of this pustule material was thawed at 4ºC in the dark and transferred to fresh BG-11 hypersaline media in a sterile culture jar and incubated for ∼2 weeks in a 12:12 h light:dark cycle to enrich for organisms that grow on cyanobacterial EPS. Following the initial incubation period, the light-grown pustule chunks were transferred into a 15 mL Falcon tube, suspended in ∼5 mL of the fresh hypersaline BG-11 medium and homogenized into a slurry using a sterile metal spatula. Triplicate sterile 50 mL glass serum bottles containing 20 mL of BG-11 medium and polysaccharide amendments were inoculated by 200 μL of pustule slurry. These cultures were capped with sterile foam plugs and kept in the dark with light shaking for 2 weeks before sampling, at which point over half of the cultures were visibly turbid. Cultures inoculated with mat slurry were grown in the dark with gentle shaking. The cultures lacking polysaccharide amendments were incubated with a 12:12 h light:dark cycle to continue to enrich for microbes that grow on cyanobacterially-produced EPS (“slurry,” see Fig. 6). After two weeks, 1.5 mL of medium was sampled from each serum bottle and the samples were stored at -20 C for ∼3 months.

Whole genomic DNA was then extracted from the samples and from ∼0.5 cm diameter chips of the thawed pustular mat material (“pustule,” see Fig. 6) using Powersoil^®^ DNA Isolation Kit (MO BIO Laboratories, Inc Carlsbad, CA, USA). DNA yield was evaluated using a Qubit 2.0 Fluorometer (Thermo Fisher Scientific, Chino, CA, USA). The extracted whole genomic DNA was submitted to the BioMicro Center Core Facility at MIT for 16s-V4 SSU amplicon (primers: U515F GTGCCAGCMGCCGCGGTAA, E786R GGACTACHVGGGTWTCTAAT) sequencing on an Illumina MiSiq sequencer with a V3 (2×150 BP) kit. Demultiplexed reads were provided by the BioMicro Core Center and further processing of the sequencing data was performed using R (v4.1.2) in the RStudio environment (Build 512). The first 20 BP of each read were trimmed to remove the non-biological fraction of the reads. Amplicon sequence variants (ASVs) were inferred using the dada2 package (V1.22.0) (83) following the guidelines of its tutorial. Due to the small overlap of the paired-end reads, we allowed merging of two reads with a minimum overlap of 3 BP. Taxonomic annotation of the ASVs was done with the RDP implementation in the dada2 package using the silva non-redundant v138.1 train set. Ampvis2 (2.7.17) (84) was subsequently used to analyze and visualize the data.

## Data availability

The Shark Bay metagenome is publicly available on the Joint Genome Institute website under the GOLD AP ID Ga0316160 (19). Amplicon sequence reads are available from the NCBI SRA under BioProject accession PRJNA826985.

## Acknowledgements

This research was funded by The Simons Foundation Collaboration on the Origins of Life (SCOL) grant number 327126 to TB and the Danish National Research Foundation grant DNRF145 to establish the Danish Center for Hadal Research, which supports CS. The APC was funded by the John V. Jarve MIT Internal Award to TB. This material is based upon work supported by the National Science Foundation Graduate Research Fellowship under Grant No. 1745302 to EC. Any opinion, findings, and conclusions or recommendations expressed in this material are those of the authors and do not necessarily reflect the views of the National Science Foundation. We thank Tania Feliz-Soto for assistance designing and preparing media, preparing colloidal chitin, and maintaining cultures in liquid media and on plates that informed the selection of polysaccharide substrates used in this study.

## Appendixes

Supplemental Data S1: S1_DegradationCAZymeActivities.xlsx

Supplemental Data S2: S2_CAZymeSubstrates.xlsx

Supplemental Data S3: S3_CAZymesInMAGsFamilyInfo.xlsx

